# Diving Behavior Reveals Humidity Sensing Ability of Water Deprived Planarians

**DOI:** 10.1101/2022.10.12.511880

**Authors:** Yu Pei, Renzhi Qian, Yuan yan, Yixuan Zhang, Liyuan Tan, Xinran Li, Chenxu Lu, Yuxuan Chen, Yuanwei Chi, Kun Hao, Zhen Xu, Guang Yang, Zilun Shao, Yuhao Wang, Kaiyuan Huang

## Abstract

Humidity sensing ability is crucial to terrestrial animals for fitting the environment. Researchers made great progress in recent study about humidity sensing mechanisms of terrestrial animals. However, it is poorly understood whether humidity sensing exists in aquatic animals. Here, we demonstrate that the aquatic planarians, one of the primitive forerunners of later animals, has the ability of humidity sensing and is capable of using the ability to perceive the water beneath itself from a drought place to seek survival. The behavior we discovered is described as diving because the worms twist its body to break away from the mucus that make them adhere to the drought place and drop into the water. The behavior is triggered by rapidly increasing humidity. This finding suggests that humidity sensing ability exists in the lower aquatic animals, and the ability might be used to seek for water when aquatic animals are facing desiccation. The finding also suggests that survival-seeking and decision-making behavior have appeared in the primitive planarian worms.

## Introduction

As a universal medium for biochemical events, water is an indispensable resource for all animals. For terrestrial animals, they are at constant risk for desiccation due to unpredictable climate change. Therefore, humidity sensing has been widely investigated in terrestrial animals for their need for a comfortable environment. Recent studies revealed detailed mechanisms of how terrestrial animals sense humidity(1-3).

In contrast to terrestrial animals, aquatic animals have much less possibility to face a situation of desiccation. However, dehydration is usually fatal to aquatic animals for they need water to respire. So, it might also be crucial for some aquatic animals to perceive the direction of water when facing an emergent situation of water depletion. Nevertheless, it is poorly understood whether this ability exists in aquatic animals.

Planarian is a kind of aquatic free-living flatworm to have first evolved a centralized brain. As a primitive forerunner of later animals, the planarians can be evolutionarily instructive for the investigation of later animals. For freely living in the natural environment, planarians have evolved various sensory abilities, including sensitivity to light(4), temperature(5), water currents(6), chemical gradients(7), vibration(8), magnetic fields(9) and electric fields(10), but its humidity sensing ability is not yet identified.

Unlike most aquatic animals who live freely in the water, planarians usually live and stick under rocks, debris and water plants in streams, ponds, and springs(11). Therefore, they are confronted with frequently falling water levels and might be lifted out of the water. So, it might be important for planarians to perceive the direction of water to seek survival under such emergent situations. Thus, we speculate that planarians have the ability of humidity sensing to carry out such tasks.

To prove this hypothesis, we established a behavioral paradigm of planarians called ‘diving’, which will be explained in detail in the result section. And then we demonstrate that the worms can perceive humidity and its increasing speed to judge the direction of the water. Our finding identified the humidity sensing ability of a kind of aquatic animal and explained what this ability is used for, which is yet not discovered in this field. This finding also suggests that survival-seeking and decision-making behavior have appeared in the primitive planarian worms and might shed light on how these abilities evolved. The finding also provides a ‘diving’ behavioral paradigm for future study.

## Results

1. Rapidly increasing humidity induces diving behavior of planarians

We established a behavioral paradigm of planarians called ‘diving’ (Fig. 1A, SI movie 1). A planarian worm is put in a petri dish and its surrounding water is wiped out. Then the petri dish is inverted onto a 250 mL beaker containing 200 mL of water. The worm will first attempt to explore around and then uplift its head. Finally, it will crawl and twist its body to break away from the mucus and drop into the water. This is more likely to be a behavior rather than a physical phenomenon.

**Figure 1.**
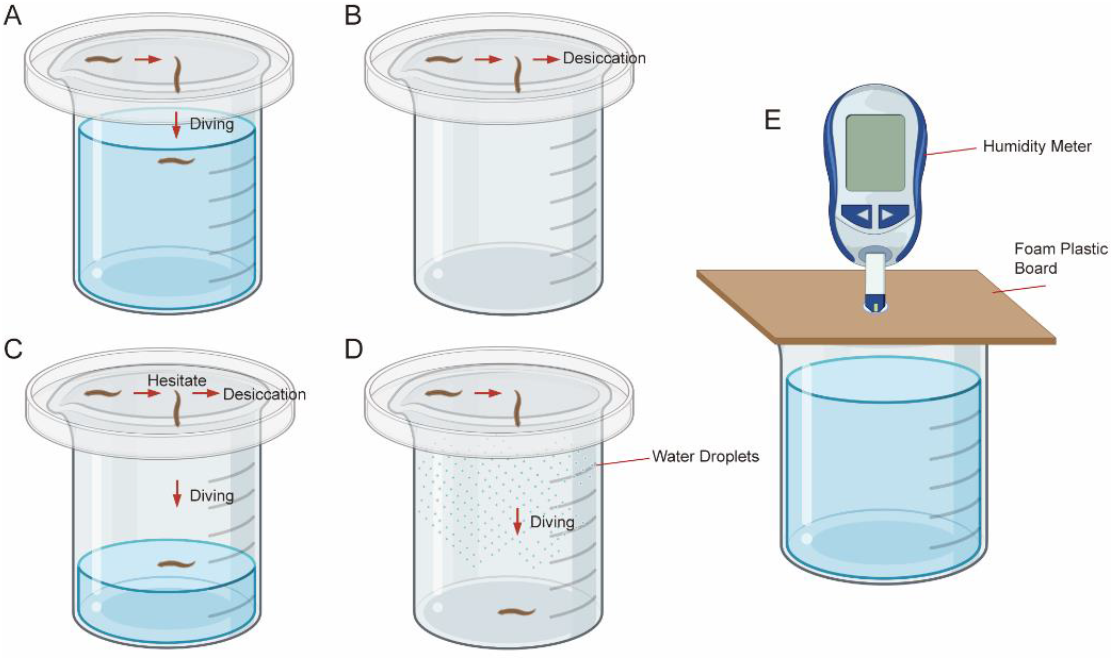
Illustration of the experiment process. (A-D) The diving experiment. (E) The relative humidity measurement.

We totally tested 20 worms through the diving paradigm and most of the worms started the diving behavior in 60 seconds, showing that they might be able to perceive the water under them. Then we tested 20 worms with a dry 250 mL beaker, all of the worms did not perform the diving behavior and finally stopped moving (Fig. 1B, SI movie 2). A worm will retract its extending head and reattach to the petri dish several times without diving. We speculate that this behavior is related to the increase in humidity. We measured the RH variation in the two processes above.

To simulate the situation of a worm in the experiment while measuring the RH variation, we embedded the humidity meter probe in the middle of a foam plastic board and then put the board on the beaker (Fig. 1E). The result reveals a rapidly increasing RH in the 250 mL beaker containing 200 mL water, which increases from 38% RH to more than 60% RH in 30 seconds. (Fig. 2A). In contrast, the RH of the dry 250 mL beaker was maintained relatively constant at 38% ±1% (Fig. 2B). The time that a worm starts the diving behavior is counted and synchronized with the RH variation. (Fig. 2A, Fig. 2F).

**Figure 2.**
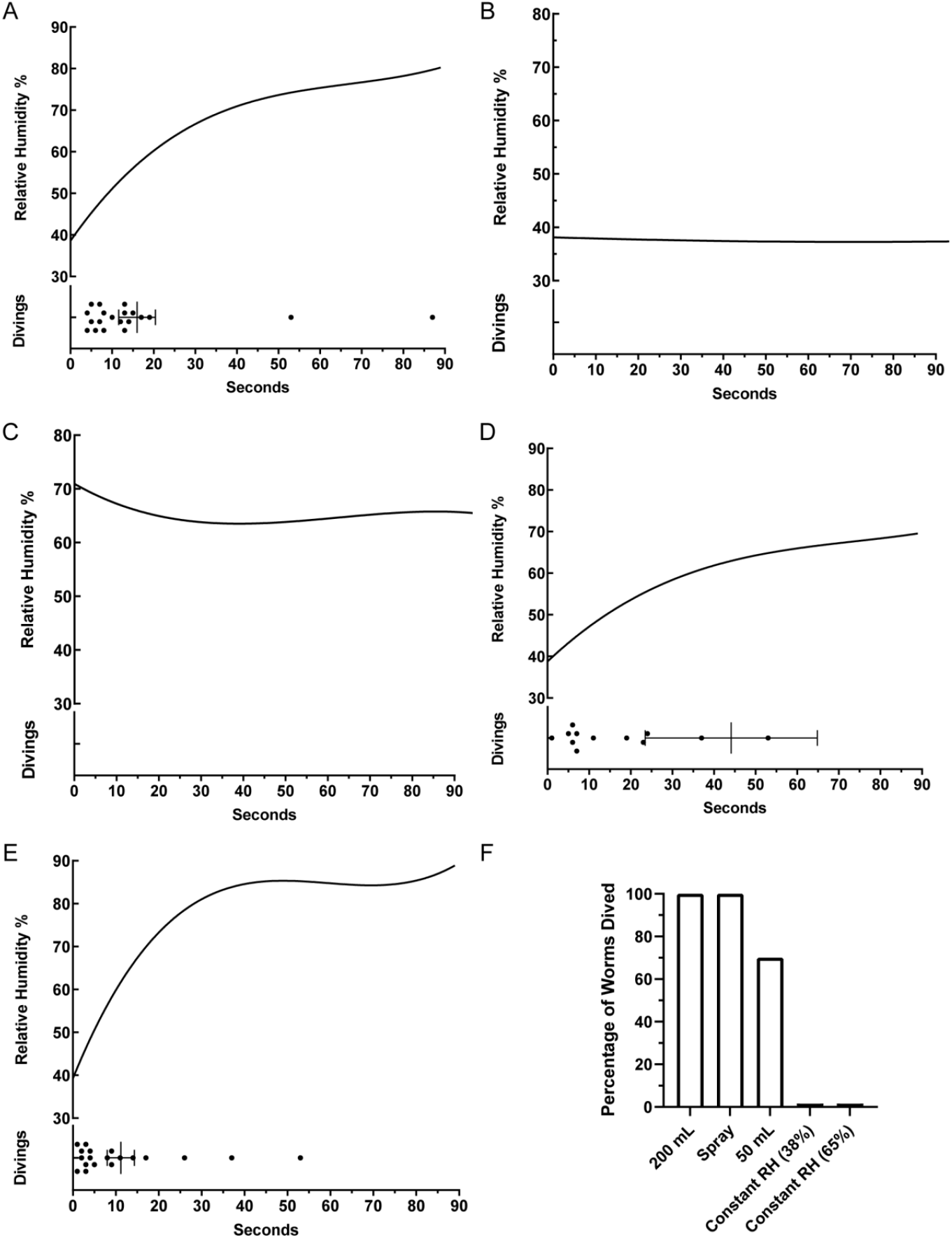
Diving behavior are induced by rapidly increasing humidity. 3 sets of RH data are used for nonlinear curve fitting in (A-E) Nonlinear regression (curve fit): polynomial (fourth order) is used to generate fitting results of RH variation. The lower panel of (A-E) shows that the time a worm starts the diving behavior synchronized with the RH variation, time data is presented as mean ± SEM. (A) Rapidly increasing humidity induces diving behavior of planarians (n=20). (B) Constant RH at around 38% cannot induce diving behavior of planarians(n=20). (C) Constant RH at around 65% cannot induce diving behavior of planarians(n=20). (D)Slower increasing humidity hinders planarians’ decision to dive (n=20, 6 worms did not dive, 2 worms used more than 90 seconds). (E) Rapidly increasing humidity can mislead planarians drop into a dry place (n=20). (F) The percentage of worms dived in each group.

We argued that the diving behavior might be induced by other factors such as temperature, density of mucus, or gravity. To control the humidity conditions, all of the diving experiment was carried out in a 38%±1% relative humidity (RH) environment if not otherwise stated. All experiments were carried out at 25±1°C to eliminate the influence of temperature. In the 250 mL group, the worms dropped at the mean time of 13 seconds and the corresponding RH is 54.1%. To exclude other factors that might interpret the diving behavior as a physical phenomenon rather than behavior, we tested 20 worms with dry 250 mL beaker at an constant RH of 65%±5% (Fig. 2C). Although the worms could struggle to crawl for a long time due to high humidity, none of the worms performed diving behavior. Hence, we conclude that the diving behavior of the worm is induced by rapidly increasing humidity rather than constant high humidity.

2. Slower increasing humidity hinders planarians’ decision to dive

To investigate whether diving behavior can be induced by slower increasing humidity, we tested 20 worms with a 250 mL beaker containing 50 mL and measured its humidity variation. (Fig. 1C, Fig. 2D). The humidity increase rate in this case is about one half of 250 mL beaker containing 200 mL of water. Surprisingly, some of the worms didn’t drop and some of the worms took about minutes to drop (Fig. 2D, Fig. 2F). We reasoned that whether to execute the diving behavior involves the worm’s decision. Slower increasing humidity makes some of the worms hesitate to drop, which further proves that only rapidly increasing humidity can solidly induce the diving behavior of planarians.

3. Rapidly increasing humidity can mislead planarians drop into a dry place

To further demonstrate the diving behavior is induced by rapidly increasing humidity, we simulated a situation of rapidly increasing humidity, yet no water is provided if the worm drops. Instead of using a large quantity of water, we sprayed water droplets on the wall of the beaker and put a piece of dry plastic to cover the bottom of the beaker (Fig. 1D). Then we tested 20 worms in this beaker and measured the humidity variation, all of the worms started to drop on the dry plastic in 60 seconds (Fig. 2E, Fig. 2F). This result confirmed the conclusion that rapidly increasing humidity induces the diving behavior of planarians.

## Discussion

We noticed that the diving of the planarian is quite different from a simple drop. A planarian will first attempt to crawl around, but soon stop crawling and extend its head. If a worm would dive, as shown by the 250 mL group (Fig. 1A, SI movie 1), it would first uplift its head (extending more than 1/2 of its body), then twist and crawl to make the tail (attached to the petri dish) move forward. As the attachment gradually decreased to a point, the planarian severely twisted its body to break away from mucus and dropped into the water eventually. If a worm would not dive, as shown by the two constant humidity groups (Fig. 1B, SI movie 2), it would also uplift its head but soon retracted its head (usually extending more than 1/2 of its body) and reattach to the petri dish. The worms might attempt to uplift its head for several times, but eventually, it will dehydrate and die on the petri dish. It seems that the planarian sensed no water below and decided not to dive. In addition, because high constant RH of 65%±5% did not trigger the diving behavior of planarians (Fig. 2C, Fig 2F), this diving behavior cannot be concluded as it is the lower density of mucus that makes worms drop and high density mucus lost moisture which prevented the drop. Above illustrates that diving is an innate behavior induced by the rapidly increasing humidity rather than a physical phenomenon.

The investigation of mechanisms of humidity sensing had been focused on terrestrial animals. Including how hygroreceptor works in insects like P. americana(12) and D. melanogaster(13), and the integration of mechano and thermo inputs of C. elegans(2) and humans(3). However, the humidity sensing of aquatic animals was hardly ever considered in previous studies. In the present study, we unveiled the ability of humidity sensing of aquatic planarians by establishing the diving behavioral paradigm, which they use to seek survival under dehydration conditions. As a kind of aquatic animal, planarians might not have hygroreceptors to directly sense water, and there is an obvious chemical barrier that would limit a planarian covered in mucus to sense the external air. Therefore, we speculate that planarians can indirectly perceive humidity through the change of mucus properties.

Our work reveals that in the diving behavioral paradigm, a worm facing dehydration has to make a quick decision whether or not to secede from the attached surface before it cannot move anymore. In this process, the worm continues to raise its head probably to sense the increasing humidity, which would accelerate the rate of evaporation. So, the judgment of the worm must be accurate to deal with such an emergent situation. As our result shows increasing rate of humidity has become a crucial indicator for the worm’s decision.

As illustrated above, the diving behavior of planarians can be classified as a decision-making and survival-seeking behavior. Being one of the first kinds of animals to have evolved a centralized brain, planarians’ behavior can provide instructions from the evolutionary perspective for investigating the behaviors of later animals. Our results demonstrate that the decision-making and survival-seeking behavior had already developed in the primitive planarian worms, which might provide a new evolutionary perspective for investigating such behaviors.

## Materials and Methods

### Planarians

A laboratory strain of *D. japonica*, originating from wild collected *D. japonica* (identified by cytochrome c oxidase subunit 1 gene) from the Cherry-Valley in Beijing Botanical Garden, Haidian district, Beijing, China in 2019. Worms are maintained in Montjuic Water(14) in the dark and fed with chicken liver twice a week. Worms are fed 2 days before experiment. The length of planarians used in the experiment varies from 1.5 cm to 2.5 cm.

### Experimental Setup

A 250mL glass beaker and a plastic petri dish is used in the experiment. The internal diameter of 250 mL beaker is 66.32 mm, the external diameter is 70.38 mm, the internal height is 94.24 mm, and the external height is 96.72 mm. Kimwipes paper towel is used to wipe water. A UT331+ humidity meter (Uni-Trend Technology (China) Co., Ltd.) is used to measure and record the RH.

### Test Procedure

A planarian is transferred to the petri dish containing water from home well by a transfer pipette. Wait until the worm sink and attach to the bottom of the petri dish. Slowly pour the water out while maintaining the worm attached to the bottom. Wipe out the water in the petri dish but not touch the worm. Then absorb water on the worm from its caudal direction until there is no water film. At this time, there is little water on the worm and no water in the petri dish. The petri dish is washed by water and wiped between each worm’s test. After the test, the animals were saved and well maintained.

### Humidity Measure Procedure

The primer of the humidity meter is embedded into a foam plastic board then cover the beaker and immediately start measuring for 90 second.

## Statistical Analysis

All data were analyzed using PRISM (GraphPad Prism 9.0.0(121)). Nonlinear regression (curve fit): polynomial (fourth order) is used to generate fitting results of RH variation.

## Acknowledgments

We wish to thank Prof. Baoqing Wang, Prof. Zhengxin Ying and associate Prof. Wei Wu for suggestions and financial support. Figure 1 is created with BioRender.com

